# Prostaglandin E₂ Reverses Myofibroblast Differentiation in Eosinophilic Esophagitis

**DOI:** 10.64898/2026.01.27.702012

**Authors:** Ryugo Teranishi, Takefumi Itami, Masaru Sasaki, Kanak V Kennedy, Yusen Zhou, Chizoba N Umeweni, Emily A Mcmillan, Archana Anandakrishnan, Reness Lee, Diya Dhakal, Hailey Golden, Grace Davis, Tatiana A Karakasheva, Mark Mahon, Bella Peterson, Heidi Winters, Nicholas Vinit, Jessi Pollack, Benjamin Wilkins, Joshua Weschler, Micheal Manfredi, Thomas E. Hamilton, Duy T. Dao, Kelly A. Whelan, Joshua Wechsler, Nancy Spinner, Emily Partridge, Amanda B Muir

## Abstract

**Background & Aims:** Unchecked inflammation in Eosinophilic esophagitis (EoE) leads to esophageal fibrosis and eventual stricture. Differentiated fibroblasts, termed myofibroblasts, are the main effector cells in fibrosis, responsible for secreting extracellular matrix proteins leading to tissue stiffness. Regulating myofibroblasts has not been explored as a therapeutic possibility in the fibrostenotic esophagus. Herein, we aim to investigate the efficacy of Prostaglandin E₂ (PGE₂) in dedifferentiation of the EoE myofibroblast.

**Methods:** We evaluated the efficacy and mechanism of myofibroblast dedifferentiation using fetal esophageal fibroblasts (FEF3), patient-derived fibroblasts, and a murine model of EoE.

**Results:** Fibrosis markers (αSMA, FN1, and COL1A1) and contractility of myofibroblasts were significantly decreased by PGE₂ via the cAMP pathway. PGE₂ treatment decreased nuclear accumulation of phospho-Smad2/3-YAP complex and induced phospho-YAP proteasomal degradation. Transcriptome analyses of FEF3 treated with TGFβ or PGE₂ revealed that the Integrin1 pathway, and specifically thrombospondin 1 (THBS-1), was significantly upregulated by TGFβ and downregulated by PGE₂, as supported by pseudo-bulk single-cell RNA-seq of EoE biopsies. THBS-1 was shown to be regulated by PGE₂ via the cAMP/YAP pathway, and its knockdown induced myofibroblasts dedifferentiation. In a murine model of EoE, Butaprost, agonist of the E-prostanoid G protein-coupled receptor 2, treatment significantly reduced the expression of THBS-1, αSMA, and FN1 along with a decrease in YAP nuclear translocation. Additionally, collagen fiber organization in the lamina propria was markedly reduced.

**Conclusion:** PGE₂ promotes dedifferentiation of myofibroblasts in EoE via the cAMP/YAP/ THBS-1 pathway. Our data suggest that PGE₂ is a promising treatment strategy for EoE with stenosis.

**What You Need to Know:** *Background and Context:* In Eosinophilic esophagitis, unchecked inflammation and tissue stiffness drives fibroblast differentiation and fibrostenosis of the esophagus, yet targeting myofibroblasts as regulators of extracellular matrix deposition in fibrostenotic disease remains clinically unexplored.

*New Findings:* Prostaglandin E₂ promotes dedifferentiation of myofibroblasts in eosinophilic esophagitis via the cAMP/YAP pathway, with Thrombospondin-1 identified as a critical YAP regulated target driving fibrostenosis.

*Limitations:* This study focused on fibroblast-specific mechanisms. The effects of PGE₂ on esophageal epithelial differentiation, barrier function, and immune cell recruitment in EoE remain to be determined.

*Clinical Research Relevance:* This study demonstrates proof-of-concept that pharmacological reversal of established fibrosis is achievable in EoE. PGE2 and its EP₂-selective agonists represent translatable therapeutic targets for fibrostenotic EoE—a patient population that remains treatment-refractory to current immunosuppressive approaches.

*Basic Research Relevance:* The cAMP/YAP/THBS-1 signaling in fibroblasts emerges as a critical therapeutic target for esophageal fibrosis. Importantly, this work demonstrates that terminally differentiated myofibroblasts retain remarkable plasticity and can dedifferentiate—challenging the paradigm that fibrosis is irreversible.

## Introduction

Eosinophilic Esophagitis (EoE) is a food allergen-triggered, cytokine-mediated chronic inflammatory disease marked by infiltration of the esophageal mucosa with eosinophils.^1^ Although clinically much attention is paid to eosinophil counts, esophageal tissue fibrosis in the lamina propria is the most severe consequence of EoE leading to recurrent food impaction, stricture and the need for dilation.^2, 3^ We previously reported that stiffness in the stromal compartment of the esophagus may contribute to myofibroblast activation and extracellular matrix (ECM) remodeling even in the absence of inflammatory cytokines.^4, 5^ These studies indicate that even in clinical remission, the esophageal microenvironment remains permissive for fibroblast activation due to persistent tissue stiffness.^4, 6^ Therefore, the development of a therapeutic intervention to interrupt continuous myofibroblast activation is critical for EoE.

The activated myofibroblast is the key effector cell in fibrosis. In particular, transforming growth factor beta (TGFβ) has been demonstrated to induce fibroblast differentiation, activation, and tissue remodeling in EoE.^3, 5^ Myofibroblasts are characterized by an increased expression of α-smooth muscle actin (αSMA) and promote fibrosis and tissue stiffness by secreting ECM proteins, such as fibronectin and collagen.^5, 7^ Current EoE therapies include proton pump inhibitors, dietary elimination, topical corticosteroids, and biologics including dupilumab,^1^ and while these are effective strategies for reducing inflammation, there is no therapy designed to mitigate the fibrosis.

Prostaglandin E_2_ (PGE_2_), a mediator of many physiological and pathological functions, is derived from cell membrane phospholipids via the cyclooxygenase pathway in the metabolism of arachidonic acid.^8, 9^ Although myofibroblasts were once considered terminally differentiated cells, some studies have shown that these cells can dedifferentiate into fibroblasts by PGE₂ in idiopathic pulmonary fibrosis (IPF).^10^ PGE₂ downregulates hallmark myofibroblast genes, including αSMA, fibronectin, and collagen I via cAMP/PKA pathway in IPF.^10^ In EoE, the narrow caliber esophagus is resistant to anti-inflammatory approaches and exploring myofibroblast dedifferentiation in EoE may provide alternative therapeutic options for treatment refractory patients.^11^

Herein, we sought to investigate the efficacy of PGE₂ for dedifferentiation of the EoE myofibroblast utilizing in vitro and in vivo models. We describe a mechanism by which PGE₂ dedifferentiates EoE myofibroblast via the cAMP/YAP/Thrombospondin-1 (THBS-1) axis.

## Materials and Methods

The materials and methods used in this study are described in detail in the Supplementary Methods.

### Human subjects and endoscopic esophageal biopsies

In accordance with the standards and guidelines of the Institutional Review Board at the Children’s Hospital of Philadelphia, esophageal biopsies were obtained from patients undergoing esophagogastroduodenoscopy as part of routine care, following informed consent. Subjects were classified based on the EoE diagnostic guidelines.^2^ The classification is provided in the Supplementary Methods, and patient demographics are provided in the Supplemental Table S1.

### Patient-derived fibroblast isolation

Patient-derived fibroblasts (PEF) were isolated from biopsy specimens as previously described with minor modifications.^5^

### Cell culture

PEF or fetal esophageal fibroblasts (FEF3) were grown at 37°C in a humidified 5% CO_2_ incubator as described previously. PEF and FEF3 culture, TGFβ–induced myofibroblast differentiation, and pharmacologic treatments (PGE₂, receptor agonists, cAMP modulators, MG-132, and Verteporfin) were carried out following the procedures described in the Supplementary Methods.

### RNA extraction and reverse transcription-quantitative polymerase chain reaction (qPCR)

RNA extraction and reverse transcription were performed as described. The used TaqMan primers are provided in Supplemental Table S2. Relative mRNA levels of each gene were normalized to *glyceraldehyde-3-phosphate dehydrogenase (GAPDH)* levels as a housekeeping control.

### Fibroblast contraction assay

Effects of PGE₂ on fibroblast contraction were determined using fibroblast contraction assays (CBA-201; Cell Biolabs, San Diego, CA) according to the manufacturer’s instructions.

### Western blotting analysis

Whole-cell lysates from cells were prepared as described previously.^12^ The primary antibodies used are provided in Supplemental Table S3. GAPDH served as a loading control.

### YAP nuclear translocation

Subcellular protein fractions were extracted from cultured FEF3 using NE PER Nuclear and Cytoplasmic Extraction Kit (78833; Thermo Scientific) according to the manufacturer’s protocol. The cytoplasmic and nuclear fractions were analyzed using Western blotting. Lamin A/C (1:1000, 2032S; Cell Signaling Technology) antibody was used for equal loading of nuclear fractions.

### Immunoprecipitation (IP)

Whole protein lysates were extracted from cultured FEF3 using Pierce™ Classic Magnetic IP/Co-IP Kit (88804; Thermo Scientific Waltham, MA) according to the manufacturer’s protocol. The IP antibodies used are provided in Supplemental Table S3.

### Immunocytochemistry

FEF3 were seeded and cultured in chamber slides (154526PK; Thermo Scientific). FEF3 were incubated with TGFβ and were treated with/without PGE₂. Cells were then incubated with the antibodies listed in Supplemental Table S3.

### Immunohistochemistry (IHC)

The IHC stain was performed on a Leica Bond RXm Autostainer (Leica Biosystems, Deer Park, IL, USA). The antibodies are listed in Supplemental Table S3. Quantification was carried out using ImageJ software (NIH, Bethesda, MD, USA).

### Masson’s trichrome staining

Masson’s trichrome staining was performed according to the protocol, which was described in the Supplementary Methods.

### Enzyme-linked immunosorbent assay (ELISA)

The Direct cAMP ELISA Kit (Enzo Life Sciences Inc., Farmingdale, NY, USA) is a competitive immunoassay that enables the quantitative determination of intracellular cAMP levels. The Human THBS-1 Quantikine ELISA Kit (DTSP10; R&D Systems) is a quantification of human THBS-1 in cell culture supernatants. Both kits were used according to manufacturer’s instructions.

### Small interfering RNA (siRNA) transfection

We performed RNA interference with siRNA directed against YAP (114602 and 135618) or THBS1 (s14099 and s14100), each consisting of 2 target-specific sequences. We used scrambled siRNA (Silencer Select Negative Control No. 1; Thermo Fisher Scientific) as nonsilencing control siRNA. The silencing efficiency was assessed by qPCR and Western blotting.

### EdU proliferation assay

The cell proliferation ability of FEF3 was carried out using a Click-iT EdU Cell Proliferation Assay (C10637; Click-iT EdU Alexa Fluor 488 Imaging Kit, Thermo Fisher Scientific). The cells were incubated with 10 µM EdU concurrently with the application of TGFβ and PGE₂, and the cell nuclei were stained with Hoechst for 30 minutes.

### mRNA sequencing and gene expression analysis

RNA sequencing was performed on FEF3 stimulated with TGFβ, treated with PGE₂, or left untreated. Differentially expressed genes were defined using a *P* value < 0.05 and predefined fold-change thresholds. Gene set enrichment analysis (GSEA; Broad Institute, Cambridge, MA) was conducted using a preranked approach to identify the top 10 enriched pathways in TGFβ-stimulated FEF3 and the top 10 depleted pathways in PGE₂-treated FEF3, selected based on the lowest FDR values from the PID database.^13, 14^ Additional experimental and computational details are provided in the Supplementary Methods.

### Fibroblast pseudo-bulk differential expression analysis

Single-cell RNA-seq data were processed using a standard workflow. Reads were aligned and quantified, followed by ambient RNA removal, quality filtering, and doublet exclusion. After dataset integration, dimensionality reduction, clustering, and cell type annotation were performed using established methods. Fibroblast clusters were subsequently extracted for pseudo-bulk RNA-seq analysis, and differentially expressed genes were identified using DESeq2 with predefined statistical thresholds. Additional computational procedures and parameter settings are provided in the Supplementary Methods.

### Murine model of EoE

BALB/c male mice (Jackson Laboratories, Bar Harbor, ME) aged 10 weeks, were procured and housed in accordance with the Institutional Animal Care and Use Committee of the Children’s Hospital of Philadelphia. Animal studies are reported in compliance with the ARRIVE guidelines. Epicutaneous sensitization of mice was performed as previously described.^15, 16^ Mice in the control and EoE groups were treated with Butaprost (4 mg/kg once daily) or Vehicle, prepared with 4% DMSO by intraperitoneal injection every day starting on day 18 for 2 weeks.

### Flow cytometry

The single-cell droplets from murine esophagi were resuspended in phosphate-buffered saline containing 4% fetal bovine serum. Dead cells were eliminated by LIVE/DEAD Fixable Aqua Dead Cell Stain Kit (1:1000, L34957; Life Technologies, Carlsbad, CA, USA). Viable cells were incubated with the antibodies listed in Supplemental Table S4. Fibroblasts were identified as CD45^-^/EpCAM^-^/DPP4^+^ PDGFRα^+^ cells.^17^ Aurora spectral analyzer (Cytek, Fremont, CA) and FlowJo software (FlowJo LLC, Ashland, OR) were used for analyses.

## Statistical analysis

Data are presented as means ± standard deviations (SD). Statistical comparisons between two groups were performed using a two-tailed Student’s t-test, and differences among multiple groups were assessed by one-way analysis of variance (ANOVA). All statistical analyses were conducted with GraphPad Prism (GraphPad Software, San Diego, CA). A *P* value less than 0.05 was considered statistically significant.

## Results

### PGE₂ decreases myofibroblast and fibrosis-associated markers in FEF3 and PEF

To determine the ability of PGE₂ to induce myofibroblast dedifferentiation, we evaluated the effects of PGE₂ treatment on the fibrosis-associated proteins and myofibroblast markers αSMA, FN1, and COL1A1 in FEF3. As depicted in the experimental scheme in Figure 1A, fibroblasts were first serum starved, followed the next day by TGFβ treatment for 24 hours, and then finally PGE₂ for 24 hours. qPCR, WB, and ICC confirmed significantly increased expression of αSMA, FN1, and COL1A1 mRNA and protein in TGFβ-induced myofibroblasts compared with untreated fibroblasts, along with decreased expression of these markers of fibroblast activation in the PGE₂ treatment groups (Figure 1B, C, and D). We next validated these findings in PEF cultures demonstrating attenuation of *ACTA2*, *FN1*, and *COL1A1* expression in TGFβ-stimulated cells (Figure 1E, Supplemental Figure S1).

**Figure 1.**
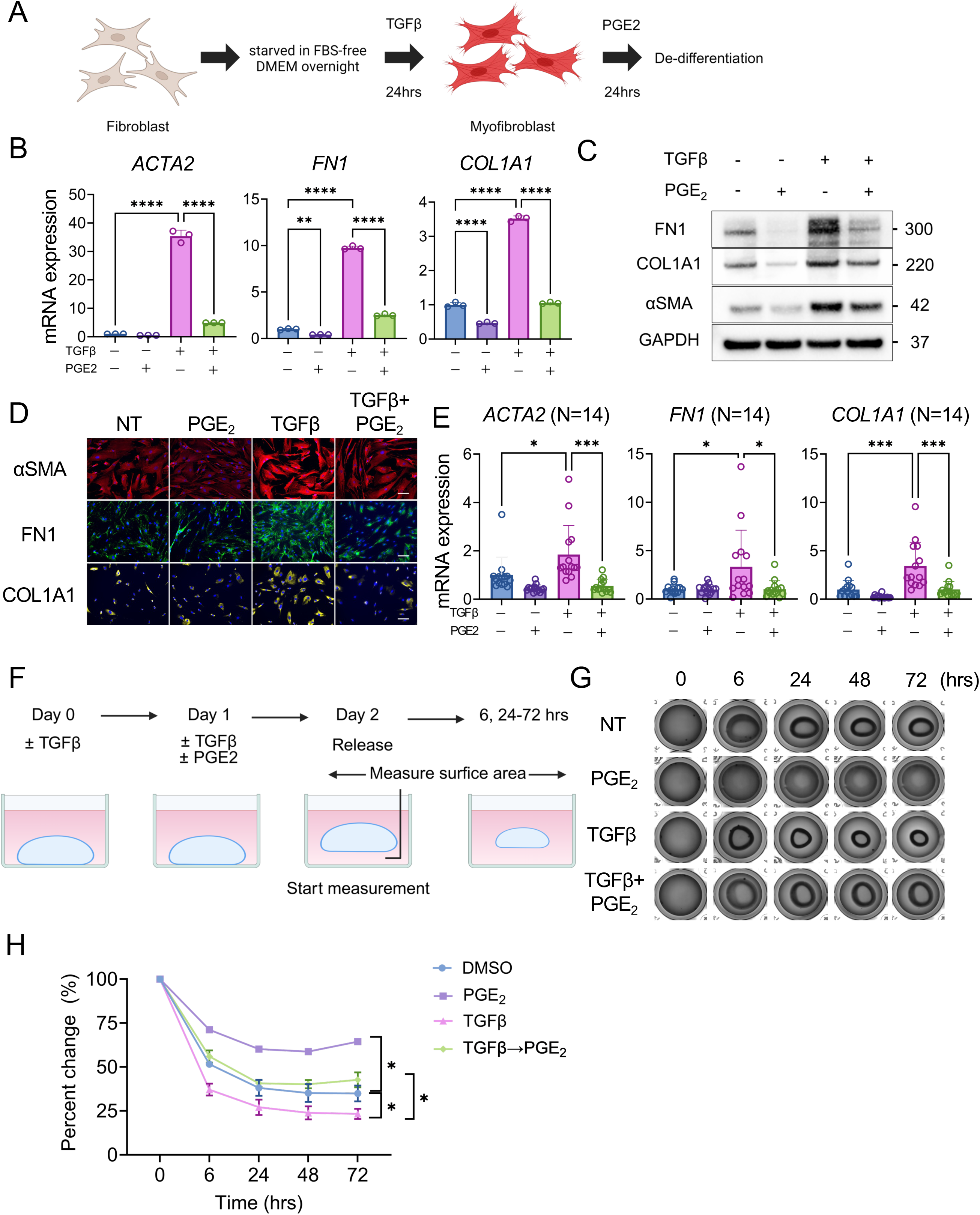
PGE₂ dedifferentiates established myofibroblasts and attenuates myofibroblast contractility. **A.** Experimental scheme depicting FEF3 and EoE patient-derived fibroblasts (PEF) differentiation with TGFβ (10 ng/mL) for 24 hours, followed by dedifferentiation with PGE₂ (1μM). **B.** Relative *ACTA2*, *FN1*, and *COL1A1* expression by qPCR in FEF3 myofibroblasts treated for 24 hours with PGE₂ (n = 3). **C.** αSMA, FN1, and COL1A1 proteins expression by western blot in myofibroblasts treated for 24 hours with PGE₂. **D.** αSMA (red), FN1 (green), and COL1A1 (yellow) were identified by immunofluorescence microscopy using anti-αSMA, FN, and COL1A1 antibodies and FITC-conjugated secondary antibody. Nuclei are stained with DAPI (blue). Scale bar 100µm. **E.** Relative *ACTA2*, *FN1*, and *COL1A1* expression by qPCR in active PEF stimulated with TGFβ with or without PGE₂ for 24 hours (n = 13) **F.** Experimental scheme depicting cell contraction assay. FEF3 (2.0×10⁶ cells/mL) were mixed with collagen solution and seeded. After collagen polymerization, FEF3 were stimulated with/without TGFβ. At 24 hours, treatment was started with/without PGE₂. At 48 hours, the gel was then mechanically detached from the plate wall. The area of each gel was measured at each time point. **G.** Representative images of collagen gels taken with the imaging system (n = 3). **H.** Change in gel area over time as measured from images. **B** and ***E.*** 1-way analysis of variance was performed for statistical analyses. * *p* < 0.05, ** *p* < 0.01, *** *p* < 0.001, **** *p* < 0.0001. **H.** A paired *t-test* was performed for statistical analyses. * *p* < 0.05

### PGE₂ decreases collagen gel contractility of myofibroblasts

Having characterized PGE₂ effects on ECM proteins and smooth muscle actin, we next sought to determine how myofibroblast function changes with PGE₂ treatment. A distinctive characteristic of myofibroblasts is their capacity for contraction. To understand if contractility is affected by fibroblast dedifferentiation, fibroblasts were seeded in collagen gels and percent contraction was measured over 3 days in the setting of TGFβ stimulation with or without PGE₂ (Figure 1F). Untreated esophageal fibroblasts demonstrated 60.4-68.0% contraction. Fibroblasts stimulated with TGFβ had significantly increased contractility (74.7-80.2%, *p* < 0.05) compared to untreated. Treatment with PGE₂ significantly decreased contraction of gels containing fibroblasts even in the setting of TGFβ (34.7-35.5%, *p* < 0.05 for PGE₂ compared with untreated, 58.9-63.8%, *p* < 0.05 for TGFβ treated with PGE₂ compared with TGFβ) (Figure 1G and H). Together, PGE₂ treatment reduced TGFβ-induced ECM expression and contractility, supporting the potential of PGE₂ induced myofibroblast dedifferentiation.

### PGE₂ employs the EP2/cAMP/PKA pathway to de-differentiate myofibroblasts

We next sought to explore the mechanism underlying PGE₂-mediated myofibroblast dedifferentiation. PGE₂ interacts with four E-prostanoid G protein-coupled receptors (EP1R to EP4R). To identify the specific receptor involved in dedifferentiation of esophageal myofibroblasts, FEF3 were serum starved prior to TGFβ and PGE₂ treatment as above.

Cultures were co-treated with selective EPR agonists, followed by analysis of fibroblast dedifferentiation. Expression of *ACTA2*, *FN1*, and *COL1A1* in myofibroblasts was strongly reduced in the presence of PGE₂ and Butaprost (EP2R agonist) (Figure 2A), indicating that PGE₂ primarily induces dedifferentiation through EP2R.

**Figure 2.**
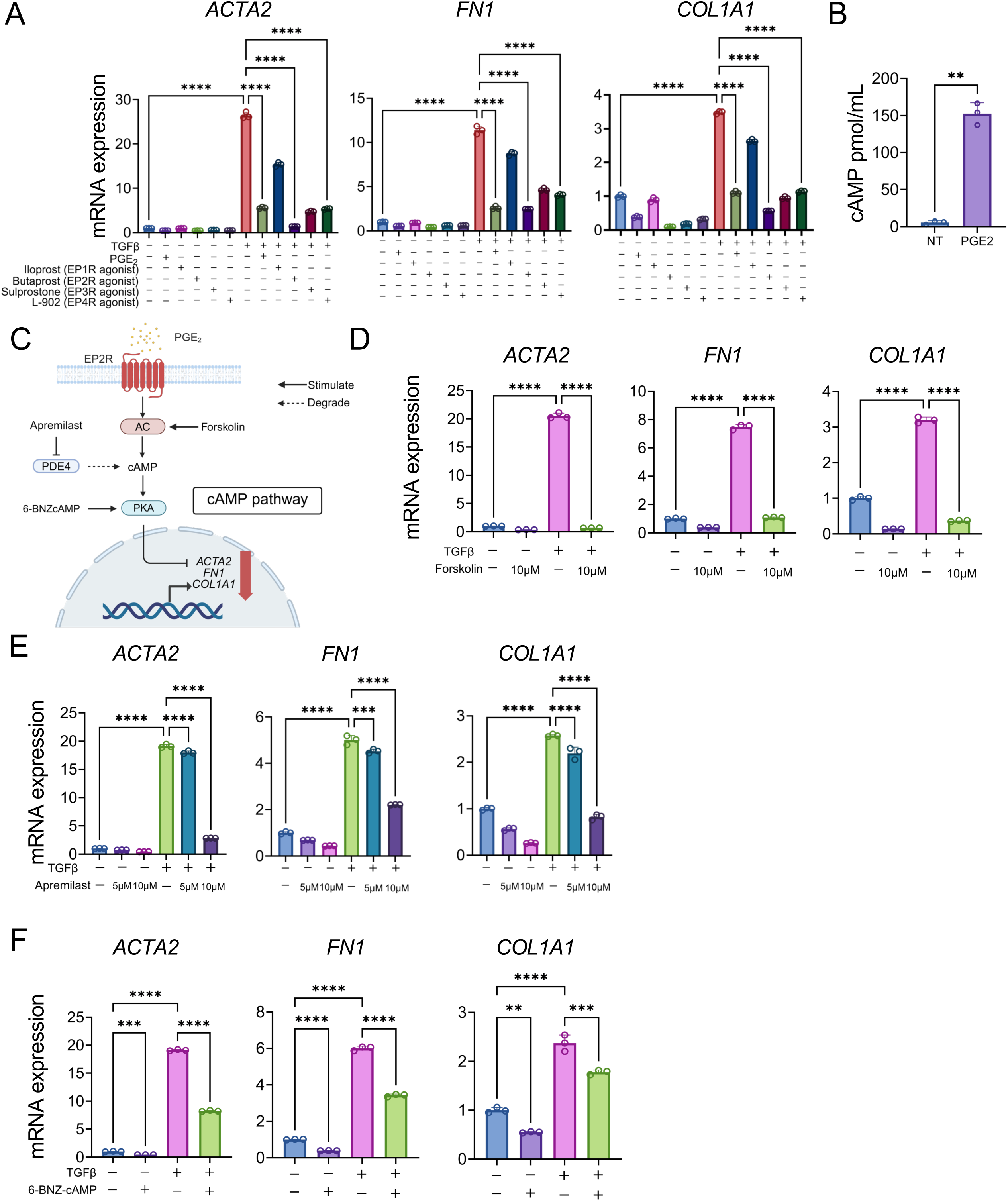
PGE₂ de-differentiates myofibroblasts via cAMP pathway. **A.** Relative *ACTA2*, *FN1*, and *COL1A1* by qPCR in FEF3 myofibroblasts treated with or without PGE₂, Iloprost (1μM; EP1R agonist), Butaprost (1μM; EP2R agonist), and Sulprostone (1µM; EP3R agonist), L-902 (1μM; EP4R agonist) (n = 3). **B.** Schematic details of the PGE₂ signaling pathway. AC, cAMP, and PKA mediate the reduction in *ACTA2*, *COL1A1*, and *FN1* elicited by PGE₂. Apremilast functions as an inhibitor of PDE4, which catalyzes the hydrolysis of cAMP. **C.** Relative intracellular cAMP by ELISA. FEF3 fibroblasts were treated with either PGE₂ for 15 minutes. **D.** Relative *ACTA2*, *FN1*, and *COL1A1* expression by qPCR in myofibroblasts treated for 24 hours with the AC activator forskolin (5 and 10µM) (n = 3). **E.** Relative *ACTA2*, *FN1*, and *COL1A1* expression by qPCR in myofibroblasts treated for 24 hours with the PDE4 inhibitor Apremilast (5 and 10 µM) (n = 3). **F.** Relative *ACTA2*, *FN1*, and *COL1A1* expression by qPCR in myofibroblasts treated for 24 hours with the PKA-specific cAMP analog 6-BNZ-cAMP (1 mM) (n = 3). Data are representative of 3 independent experiments and expressed as means ± SDs. **A, C, E,** and **F.** 1-way analysis of variance was performed for statistical analyses. **D.** Paired *t-test* was performed for statistical analyses. ** *p* < 0.01 *** *p* < 0.001, **** *p* < 0.0001. adenylate cyclase; AC, protein kinase A; PKA, Phosphodiesterase; PDE

PGE₂ is known to activate the cAMP pathway through EP2R (Figure 2B).^9^ Indeed, in esophageal fibroblasts, ELISA showed that treatment with PGE₂ significantly increased intracellular cAMP levels after 15 min (Figure 2C). Upon activation of EP2 by PGE₂ or Butaprost, it has been shown that adenylyl cyclase (AC) is stimulated, increasing intracellular cAMP levels and subsequently activating protein kinase A (PKA) (Figure 2B).^18^ We investigated the capacity of the cAMP pathway components to induce dedifferentiation in esophageal myofibroblasts. Forskolin, a direct activator of AC, reduced levels of *ACTA2*, *FN1*, and *COL1A1* (Figure 2D) in TGFβ activated fibroblasts, similar to the trend observed with PGE₂ treatment. In addition, Apremilast, a PDE4 inhibitor which serves to increase intracellular cAMP, and 6-BNZ cAMP, a cAMP analog that selectively activates PKA, induced myofibroblast dedifferentiation with decreased *ACTA2*, *FN1*, and *COL1A1* (Figure 2E and F). This suggests that PGE2 drives myofibroblast dedifferentiation primarily through upregulation of the cAMP pathway through the receptor EP2.

### YAP plays a crucial role in the dedifferentiation of myofibroblasts

TGFβ stimulation induces activation of fibroblasts in organs such as the heart, lungs, and liver through the nuclear accumulation of phosphorylated Smad2/3 (pSmad2/3), where it forms a complex with YAP.^19–21^ Thus, we investigated the role of YAP in EoE fibrosis. We performed IHC for YAP in esophageal biopsy specimens with adequate representation of lamina propria based on H&E staining. Overall expression of YAP as well as nuclear localization of YAP in the lamina propria was significantly higher in active EoE vs control subjects. Biopsies from patients with EoE in remission demonstrated YAP expression and localization similar to control subjects (Figure 3A). To determine if YAP plays a role in fibroblast activation, we utilized siRNA knockdown of YAP in FEF3. We found that YAP knockdown decreased expression of αSMA, FN1, and COL1A1 by Western blot in TGFβ stimulated myofibroblasts (Figure 3B and Supplemental Figure S2). Furthermore, Verteporfin, a YAP inhibitor, decreased *ACTA2*, *FN1*, and *COL1A1* expression in the myofibroblasts (Supplemental Figure S3). This demonstrates that YAP is essential for myofibroblast activation in the EoE esophagus.

**Figure 3.**
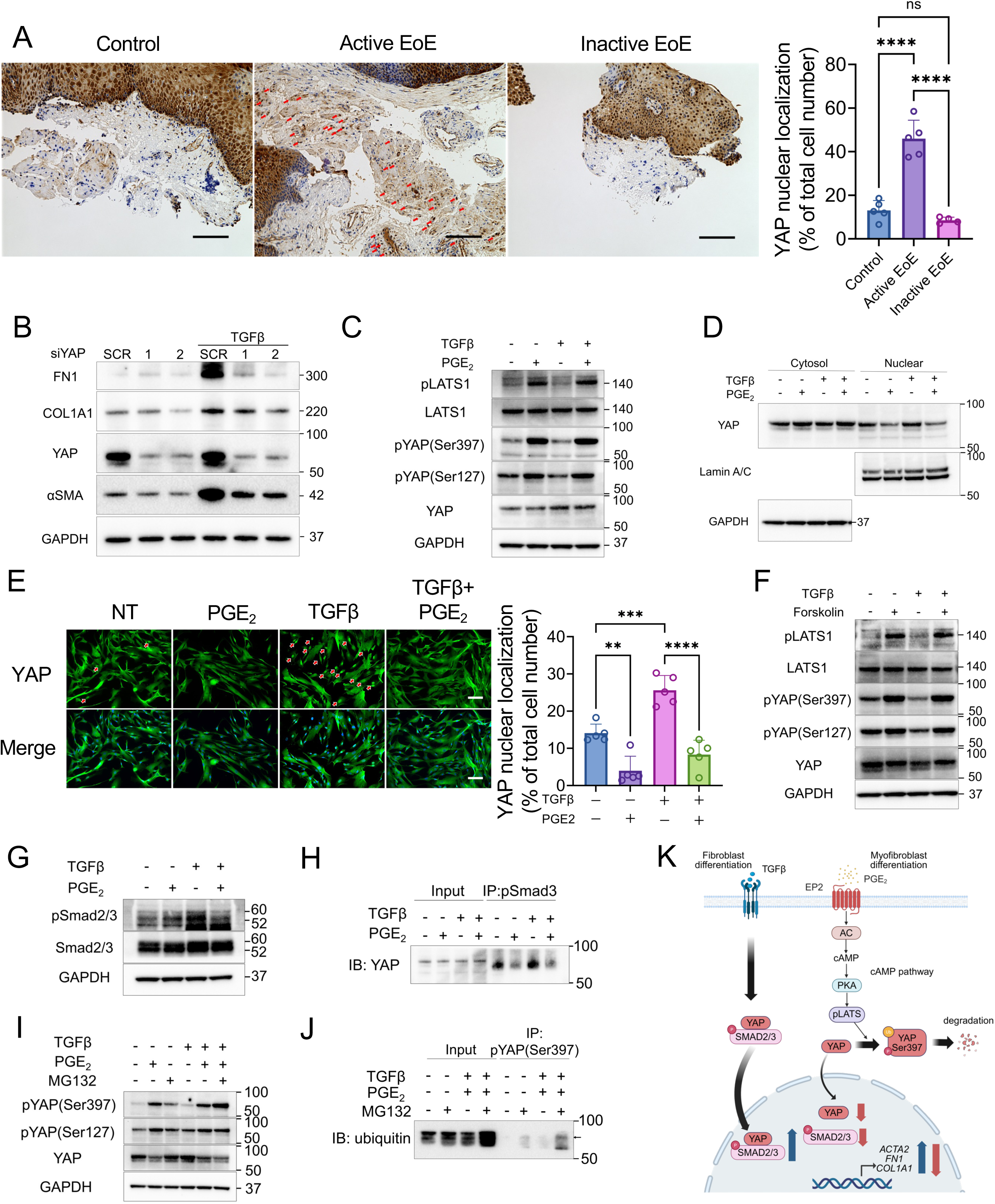
PGE₂ dedifferentiates myofibroblasts via attenuation of the LATS–YAP pathway.Phosphorylated YAP is degraded by proteasome via the cAMP pathway. A. Representative images of immunohistochemistry of YAP in human biopsy tissues. Nuclear YAP (Red allow). Quantifying the percentage of cells with YAP in the nucleus per high-power fields (HPFs). Data were obtained from five subjects in the active EoE and normal groups and from three subjects in the remission group. For each subject, five randomly selected HPFs were analyzed, and the mean value per subject was calculated. Scale bar 100µm. **B.** Transfection with siRNA against YAP in FEF3. Scramble siRNA-transfected FEF3 were used as a control. Twenty-four hours after transfection, FEF3 was stimulated with TGFβ. Relative YAP, αSMA, FN1, and COL1A1 protein expression by Western blot. **C.** Representative Western blots showing phosphorylation of YAP, phosphorylation of LATS1, and their total protein expression were determined 30 minutes after treatment with TGFβ (10 ng/mL) with or without PGE₂ (1 µM), respectively. **D.** Nuclear translocation of YAP by Western blot. FEF3 were treated with TGFβ with or without PGE₂ for 30 minutes, and nuclear and cytosolic fractions were separated. **E.** Representative images of immunofluorescence of YAP (green) in FEF3. Nuclei are stained with DAPI (blue). Nuclear YAP (Red allow). Quantifying the percentage of cells with YAP in the nucleus per HPFs. For each condition, five randomly selected HPFs were analyzed, and the mean value per condition was calculated. Scale bar 100µm. **F.** Representative Western blot showing phosphorylation of YAP, phosphorylation of LATS1, and their total protein expression were determined 30 minutes after treatment with TGFβ (10 ng/mL) with or without Forskolin (1 µM), respectively. **G.** Relative phosphorylation and total Smad2/3 protein expression by Western blot. **H.** Following immunoprecipitation of phosphorylated Smad3, an immunoblot of YAP was conducted. FEF3 were treated with TGFβ with or without PGE₂ for 30 minutes. **I.** FEF3 were pretreated with or without MG132 (10 µM) for 6 hours, exposed to TGFβ with or without PGE₂ for 30 minutes, and subjected to Western blot analysis of phosphorylation and total YAP. **J.** Following immunoprecipitation of phosphorylated YAP(Ser397), an immunoblot of ubiquitin was conducted. FEF3 were pretreated with or without MG132 for 6 hours, and exposed to TGFβ with or without PGE₂ for 30 minutes. **K.** Schematic details of the myofibroblast dedifferentiation via canonical TGFβ and YAP signaling pathways. **E.** 1-way analysis of variance was performed for statistical analyses. ** *p* < 0.01 *** *p* < 0.001, **** *p* < 0.0001, ns not significant.

In addition to its role in the TGFβ pathway, YAP is also regulated by cAMP through the LATS kinase. Specifically, LATS activity is regulated through phosphorylation by PKA.

Phosphorylation of YAP by LATS on Ser127 and Ser397 prevents its nuclear localization and leads to its proteasomal degradation.^22, 23^ To investigate the role for PGE₂ in YAP inhibition, we examined phosphorylation of YAP and LATS in esophageal fibroblasts and myofibroblasts. We observed no difference in total YAP or phosphorylated YAP (pYAP) with TGFβ stimulation. However, 30 minutes after the addition of PGE₂, phosphorylation of LATS as well as YAP (pYAP Ser397/Ser127) was increased (Figure 3C). We next examined YAP nuclear translocation in fibroblast differentiation and observed that TGFβ promoted nuclear translocation of YAP from the cytosol, suggesting its activation (Figure 3D and E). Conversely, PGE₂ significantly inhibited both basal and TGFβ-mediated YAP translocation (Figure 3D and E). This suggests that PGE₂ induces myofibroblast dedifferentiation by preventing translocation of YAP to the nucleus.

We next sought to validate that PGE₂ regulates YAP via the cAMP pathway. We examined changes in pYAP and phosphorylated LATS using forskolin and 6BNZ-cAMP. Similar to PGE₂, forskolin and 6BNZ-cAMP increased YAP and LATS phosphorylation (Figure 3F and Supplemental Figure S4). This suggests that myofibroblasts de-differentiate in response to PGE₂ by an increase in cAMP and PKA activity which leads directly to loss of YAP nuclear localization, presumably preventing YAP functioning as a regulator of a transcriptional program.

### PGE₂ decreases YAP and phospho-Smad2/3 complex and induces phospho-YAP degradation

The canonical TGFβ pathway induces fibroblast differentiation via Smad2/3 phosphorylation in EoE.^4, 5^ YAP has been shown to interact with Smad complex sequestering it to the nucleus and driving fibrogenesis.^20, 24^ However, the role of YAP in TGFβ signaling in esophageal fibroblasts is unknown. We investigated the mechanism by which PGE₂ attenuates the TGFβ pathway via the inhibition of YAP. As expected, pSmad2/3 was increased in esophageal myofibroblasts compared to fibroblasts, and it was decreased by PGE₂ (Figure 3G). To investigate the effect of PGE₂ on the YAP-pSmad2/3 complex, we performed immunoprecipitation with phospho-Smad3. Immunoblotting for YAP revealed an increase in the YAP-pSmad3 complex following stimulation with TGFβ and a decrease in the complex following treatment with PGE₂ (Figure 3H). The result of the negative control using normal IgG is shown in Supplemental Figure S5. We next investigated the mechanism by which YAP is degraded after phosphorylation. Prior to stimulation with TGFβ and PGE₂, fibroblasts were pretreated with MG132, a proteasome inhibitor, for 6 hours under basal conditions. We found increased pYAP(Ser397) in MG132 treated myofibroblasts compared to myofibroblasts treated with only TGFβ and PGE₂ (Figure 3I), indicating that pYAP(Ser397) is degraded by the proteasome. The result of the negative control using normal IgG is shown in Supplemental Figure S6. To further confirm proteasomal degradation, cell lysates were immunoprecipitated using pYAP(Ser397) antibody, and the immunoblots were probed for ubiquitin (Figure 3J). The result demonstrated an increase in ubiquitinated proteins in the MG132 pretreated group. Taken together, these results indicate that the YAP-pSmad2/3 complex mediates the canonical TGFβ pathway, while PGE2 signaling mediates phosphorylation of YAP and its subsequent ubiquitylation and degradation by proteasomes (Figure 3K).

### PGE2 regulates myofibroblast differentiation by suppressing TGFβ-activated Integrin 1 pathway

To understand the mechanism by which PGE₂ regulates myofibroblast dedifferentiation, we performed RNA sequencing on FEF3 treated with TGFβ or PGE₂, and also conducted pseudo-bulk sequencing of fibroblasts isolated from EoE subjects with active disease and in remission. (Figure 4A). GSEA on the PID revealed that stimulation of FEF3 with TGFβ resulted in an enrichment in the Integrin 1 pathway (9 DEGs involved) (*p* < 0.05 and log_2_(fold change) > 1.5) (Figure 4B). Stimulation of fibroblasts with PGE_2_ resulted in enrichment of the Integrin 1 pathway (5 DEGs involved) (*p* < 0.05 and log_2_(fold change) < - 1.5) (Figure 4C). When comparing fibroblasts from biopsies of patients with active EoE to patients with remission, in the active EoE fibroblast, PID_ Integrin1_ pathway was also enriched (16 DEGs involved) (*p* < 0.05 and log_2_(fold change) > 1.5) (Figure 4D). These results suggest that the Integrin1 pathway plays a crucial role in myofibroblast dedifferentiation. Therefore, we explored genes involved in dedifferentiation within the Integrin 1 pathway. In those 3 groups, only *THBS1* and *FN1* were present in each group (Figure 4E). *THBS1* demonstrated high normalized counts and was significantly upregulated or downregulated in fibroblasts stimulated with TGFβ or PGE₂, respectively (Figure 4F). Therefore, we explored genes involved in dedifferentiation within the Integrin 1 pathway. In those 3 groups, only *THBS1* and *FN1* were present in each group (Figure 4E). *THBS1* demonstrated high normalized counts and was significantly upregulated or downregulated in fibroblasts stimulated with TGFβ or PGE₂, respectively (Figure 4F). *THBS1* gene encodes THBS-1 protein, known to interact with a variety of integrins.^25^ Therefore, we postulated that THBS-1 could be critical in PGE₂-induced myofibroblast dedifferentiation.

**Figure 4.**
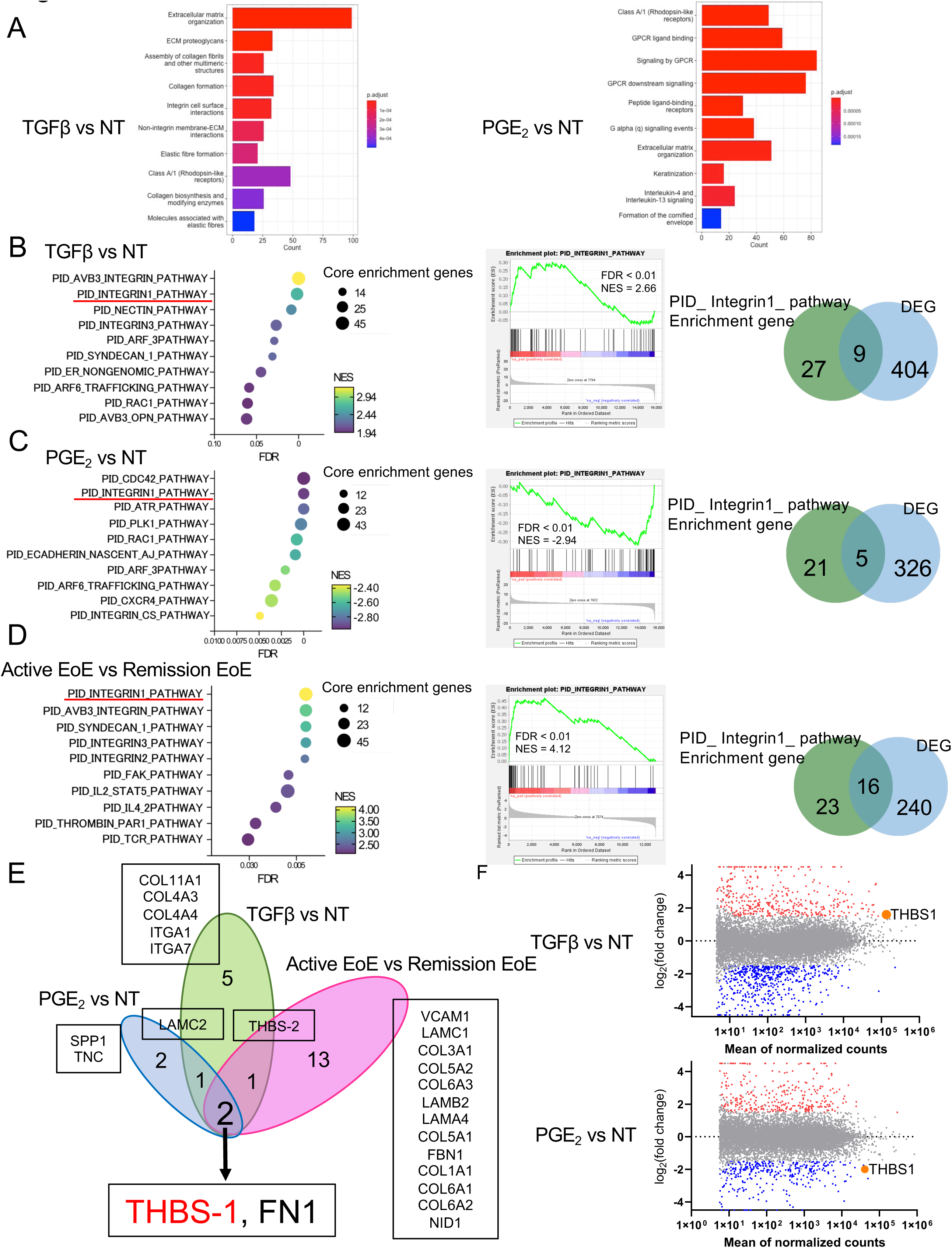
RNA-seq of FEF3 stimulated with TGFβ or treated with PGE₂, and pseudo-bulk EoE fibroblast in active and remission EoE. **A.** Experimental schematic of bulk RNA-seq analysis of FEF3 stimulated with TGFβ or treated with PGE₂, and pseudo-bulk sequencing of fibroblasts isolated from subjects with active EoE or remission. **B. C. D.** Top 10 enriched terms in TGFβ stimulated FEF3 and depleted terms in PGE₂ treatment FEF3. And Top 10 enriched terms in fibroblasts from active EoE patients compared with remission EoE patients. Gene Set Enrichment Analysis for Integrin1 pathway in the PID based on each condition. Venn diagrams depicting enrichment gene in PID_ Integrin1 pathway and the number of DEGs. The threshold for DEGs set by log2 fold change –1.5 or 1.5 and *p* < 0.05. All data sets were analyzed on Gene Set Enrichment Analysis based on the PID. Dot size and color represent the number of core enrichment genes and the normalized enrichment score (NES) for the pathway. FDR, false discovery rate. **E.** Venn diagrams depicting overlapping genes in each condition (Figure 4 B, C, and D). **F.** MA plots based on RNA sequencing data. The x-axis is the mean of the normalized counts and the y-axis is the log2(fold change). DEGs with *p* < 0.05 and log2(fold change) > 1.5 were plotted in red, and DEGs with *p* < 0.05 and log2(fold change) <-1.5 were plotted in blue. THBS-1 was plotted as orange.

### THBS-1 is crucial in fibroblast differentiation

We validated increased expression in *THBS-1* by qPCR in FEF3 and PEF in the setting of TGFβ (Figure 5A and 5B). PGE₂ treatment decreased the expression of *THBS-1* with or without TGFβ (Figure 5A and 5B). Interestingly, we found in PEF that active EoE fibroblasts (n = 7) expressed significantly more *THBS1* than remission (n = 7) (Figure 5C). THBS-1 was expressed robustly in active EoE subjects and decreased in patients in remission and controls. (Figure 5D). THBS-1 is secreted by fibroblasts. To evaluate the effect of PGE₂ on THBS-1 secretion, we tested the conditioned media from PGE₂-treated fibroblasts and myofibroblasts by ELISA. We observed that PGE₂ treatment significantly decreased extracellular THBS-1 (Figure 5E). Next, to investigate the *YAP*- *THBS-1 axis*, we performed *YAP* knockdown using *YAP*-specific siRNA. siYAP decreased *THBS-1* expression in myofibroblasts (Figure 5F). Consistent with siYAP, verteporfin also decreased expression of *THBS-1* in myofibroblasts (Figure 5G). To confirm a role for THBS-1 in fibroblast gene expression, we knocked down expression using *THBS1*-specific siRNA and investigated the expression of the myofibroblast markers αSMA, FN1, and COL1A1. siTHBS-1 decreased FN1 and COL1A1 expression but not αSMA (Figure. 5H). Interestingly, stimulation with THBS-1 increased fibroblast proliferation, and siTHBS-1 decreased fibroblast proliferation despite stimulation with TGFβ (Figure 5I and 5J). These findings indicate that THBS-1 is crucial for fibroblast differentiation and proliferation.

**Figure 5.**
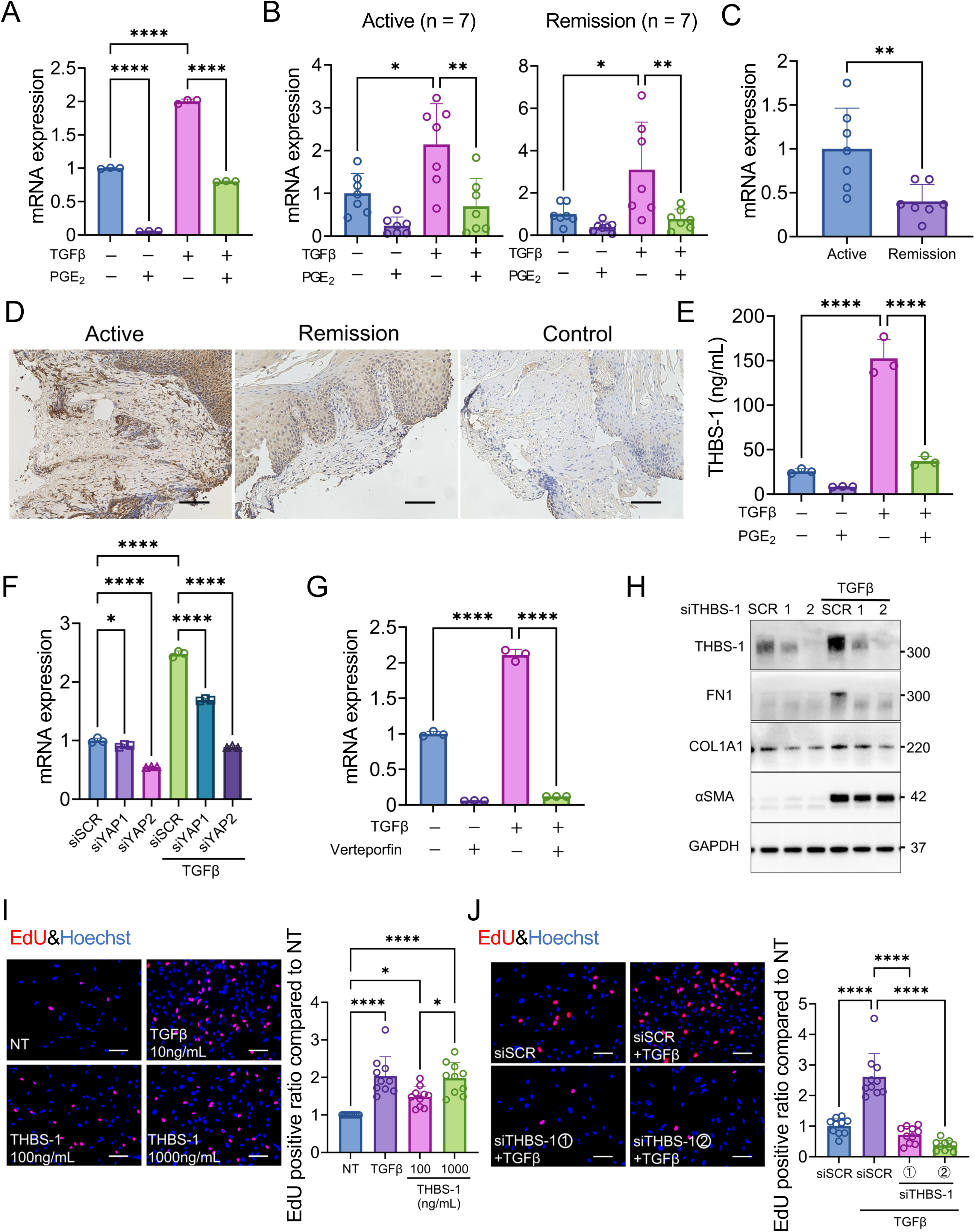
THBS-1 is subject to YAP regulation and plays a crucial role in the differentiation and proliferation of fibroblasts. **A.** Relative *THBS-1* expression by qPCR in FEF3 myofibroblasts treated for 24 hours with PGE₂ (1 μM) (n = 3). **B.** Relative *THBS-1* expression by qPCR in fibroblasts from active and remission EoE patients stimulated with TGFβ (10 ng/mL) with or without PGE₂ for 24 hours (n = 7). **C.** Relative *THBS-1* expression by qPCR in fibroblasts from active and remission EoE patients (n = 7) **D.** Representative images of immunohistochemistry of THBS-1 in human biopsy tissues. Scale bar 100µm. **E.** Relative THBS-1 protein in supernatant by ELISA. FEF3 were stimulated with TGFβ and treated with or without PGE₂ for 15 minutes. **F.** Transfection with siRNA against YAP in FEF3. Scramble siRNA-transfected FEF3 were used as a control. Twenty-four hours after transfection, fibroblasts were stimulated with TGFβ. Relative *THBS-1* expression by qPCR (n = 3). **G.** Relative THBS-1 expression in FEF3 myofibroblasts treated with verteporfin by qPCR (n = 3). **H.** Transfection with siRNA against THBS-1 in FEF3. Scramble siRNA-transfected FEF3 were used as a control. Twenty-four hours after transfection, fibroblasts were stimulated with TGFβ. Relative THBS-1, αSMA, FN1, and COL1A1 proteins expression by western blot. **I.** The EdU staining is used to detect cell proliferation. The cell nuclei are stained blue (DAPI), and the EdU+ nuclei are stained red. Quantifying the percentage of EdU+ nuclei in the nucleus per HPFs. For each condition, randomly selected HPFs (n = 10) were analyzed, and the mean value per condition was calculated. FEF3 stimulated with TGFβ and THBS-1 (100 and 1000 ng/mL). Scale bar 100µm. **J.** The EdU staining is used to detect cell proliferation. Transfection with siRNA against THBS-1 in FEF3. The cell nuclei are stained blue (DAPI), and the EdU+ nuclei are stained red. Quantifying the percentage of EdU+ nuclei in the nucleus per HPFs. For each condition, randomly selected HPFs (n = 10) were analyzed, and the mean value per condition was calculated. Scale bar 100µm. All data were representative of 3 independent experiments and expressed as means ± SDs. **A, B, C, D, E, F,** and **G.** 1-way analysis of variance was performed for statistical analyses. * *p* < 0.05, ** *p* < 0.01, *** *p* < 0.001, **** *p* < 0.0001.

### Butaprost induces myofibroblast dedifferentiation in a murine model of EoE

To establish the effects of PGE₂ in murine EoE, we employed the selective EP2R agonist Butaprost, which exhibited comparable or superior efficacy to PGE₂ in vitro (Figures 2A). Butaprost administration started after the induction of EoE to assess whether it could induce EoE myofibroblast dedifferentiation (Figure 6A). We found that mice with experimental EoE treated with Butaprost had decreased collagen deposition compared to vehicle-treated controls (Figures 6B and 6C). THBS-1 expression and YAP nuclear localization in the esophageal lamina propria were also reduced in the Butaprost group (Figures 6B and 6C). Flow cytometry revealed decreased THBS-1 expression in the Butaprost-treated group. Expression of the myofibroblast markers αSMA and FN1 was likewise reduced in Butaprost-treated mice (Figures 6D and 6E). The gating strategy for flow cytometry is provided in Supplemental Figure S7.

**Figure 6.**
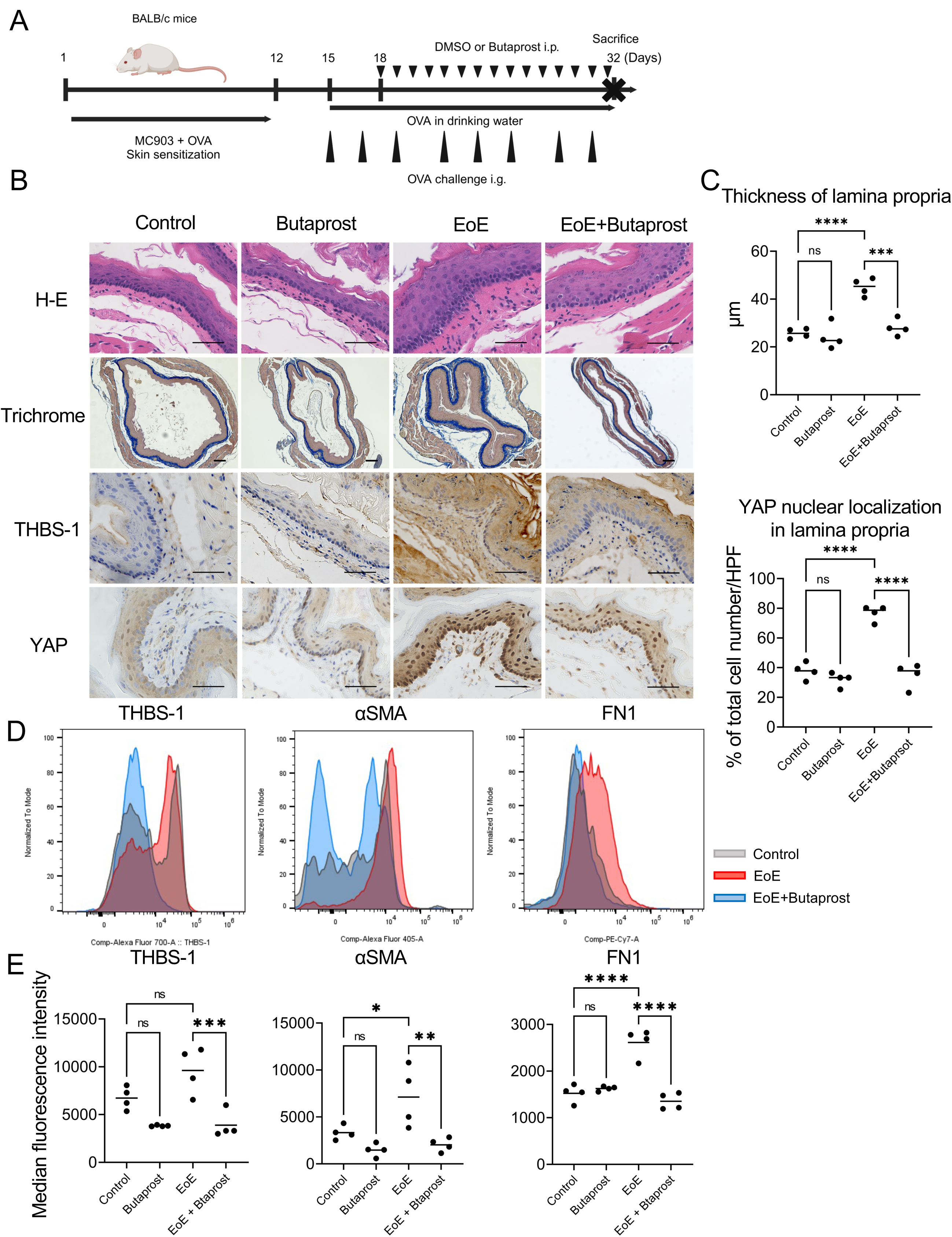
Butaprost de-differentiates myofibroblasts and collagen deposition via YAP/THBS-1 axis. **A.** Schematic of murine EoE model and Butaprost treatment. **B.** Representative images of hematoxylin and eosin (H&E), trichrome staining, immunohistochemistry for THBS-1 and YAP of the murine esophagus. Scale bar, 50 μm for H&E, THBS-1, and YAP, 100μm for trichrome staining. **C.** Thickness (μm) of lamina propria and number of nuclear YAP-stained cells per high-power field. Thickness was measured in 10 randomly selected high-power fields (HPFs), and YAP staining was analyzed in five randomly selected HPFs. Data were obtained from four mice per group. Group means were calculated from individual mouse averages. **D.** Representative flow cytometry histogram of THBS-1, αSMA, and FN1 in the murine esophagi. **E.** Quantification of THBS-1, αSMA, and FN1 expression in the murine esophagi as measured by flow cytometry (n = 4). Data are indicated as means ± SDs. One-way analysis of variance (C, D, and E) was utilized for statistics. * *p* <0.05, ** *p* <0.01, *** *p* <0.001, **** *p* <0.0001. ns not significant

## Discussion

EoE is characterized as a progressive fibrostenotic disease with approximately 10% of EoE patients developing an extremely narrow-caliber esophagus.^11, 26–28^ Recent studies have shown that decreased distensibility and fibrostenosis of the esophagus in EoE are associated with esophageal dysmotility.^29–31^ This underscores the need to better understand the often neglected remodeling that is going on below the surface epithelium. Current therapeutic approaches target the immune infiltrate, and there is no fibroblast directed therapy to decrease the effects of the activated myofibroblasts and prevent ongoing esophageal stiffening. Herein, we show that treatment with PGE₂ leads to myofibroblast dedifferentiation. PGE₂ induces YAP phosphorylation and proteasomal degradation via the cAMP pathway and thus leads to decreased expression of THBS-1 and myofibroblast activation. Furthermore, decreased THBS-1 alone reduces ECM proteins FN1 and COL1A1 as well as fibroblast proliferation. Taken together, PGE₂ induces myofibroblast dedifferentiation and decreases proliferation via the cAMP/YAP/THBS-1 axis (Figure 7).

**Figure 7.**
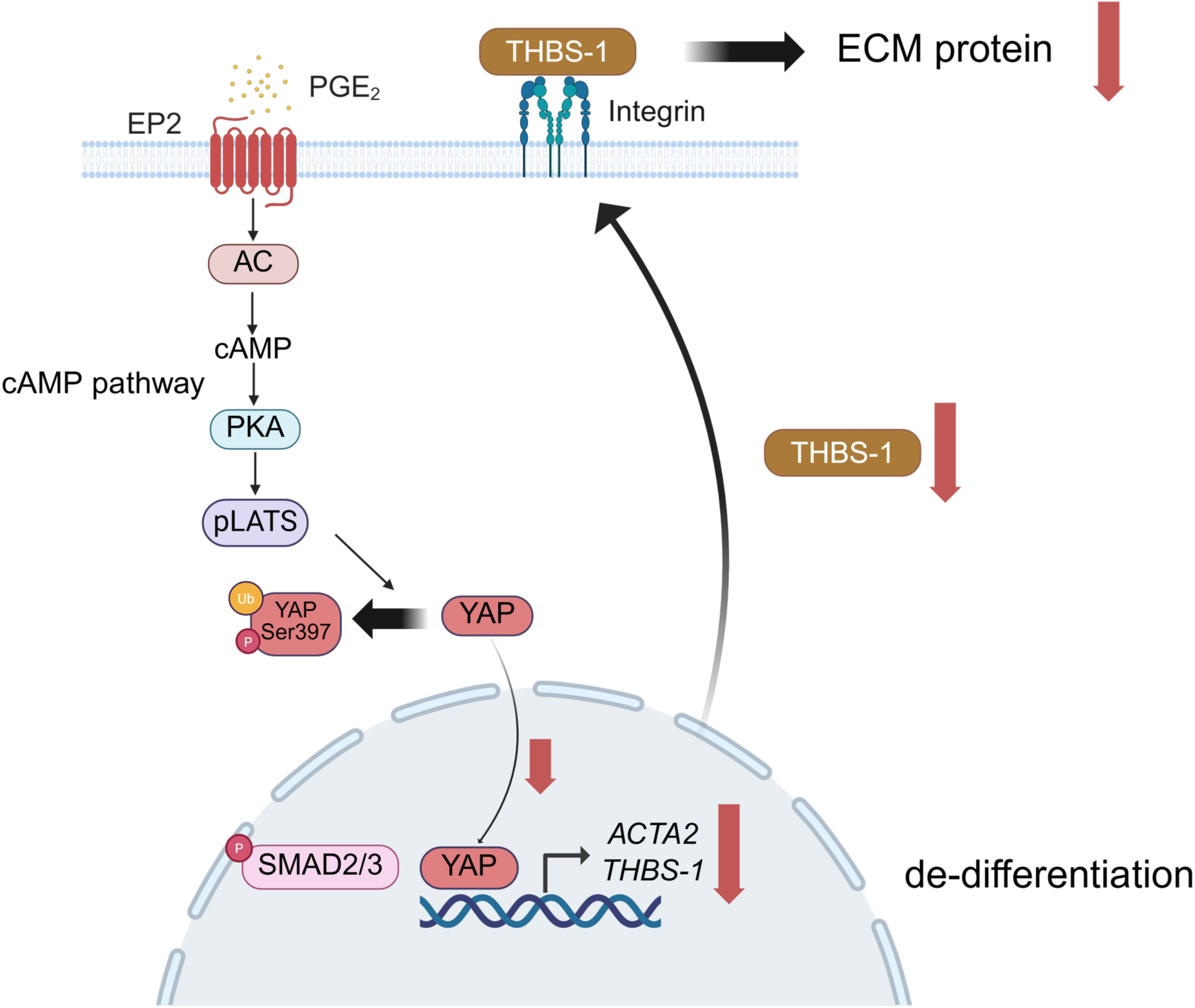
Models of myofibroblast differentiation via cAMP/YAP/THBS-1 by PGE₂. PGE₂ induces YAP phosphorylation and degradation by proteasome via the cAMP pathway. YAP downregulation induced decreasing αSMA and THBS-1. Furthermore, decreasing THBS-1 induces downregulating ECM proteins such as FN1 and COL1A1 and fibroblast proliferation.

These results present a potential way to interrupt continuous myofibroblast activation in patients with EoE.

PGE₂ binds to its cognate cell surface receptors, EP1, EP2, EP3, and EP4. We have shown that in esophageal fibroblasts PGE₂ prevents TGFβ-induced fibroblast differentiation mainly through activation of the EP2 receptor, in agreement with previous reports in IPF.^10^ We now report that modulation of intracellular cAMP mitigates fibroblast differentiation in the esophagus. PGE₂ is used clinically for evacuation of uterine contents and induction of labor.^32^ Harnessing this pathway may offer a new approach to mitigating fibrostenosis in EoE, which could be used as an add-on to current anti-inflammatory approaches to treatment.

YAP, the pivotal mediator of mechanotransduction, is a transcriptional cofactor that shuttles between the nucleus and cytoplasm.^33, 34^ Nuclear localization of YAP may establish a feed-forward loop that further enhances ECM proteins in response to a rigid substrate like ECM. Rawson et al. reported increased numbers of smooth muscle cells with nuclear-localized YAP in EoE, however, nothing is known about its role in the EoE fibroblast.^33^ YAP may represent a critical target for regulating myofibroblasts in EoE. We demonstrated that PGE₂ attenuates YAP-induced fibrogenesis by promoting LATS phosphorylation and subsequent YAP degradation, thereby inhibiting canonical TGFβ signaling. Given the reported crosstalk between YAP and Smad-mediated transcription,^19–21^ inhibition by PGE₂ may disrupt a cooperative profibrotic signaling axis. This disruption could extend to other downstream targets of TGFβ that modify the ECM and its stiffness, such as lysyl oxidase.^4^

The ECM provides not only a simple scaffold but also a repository for a variety of different growth factors and matricellular proteins.^35^ THBS-1 is a member of a family of calcium-binding glycoproteins and a matricellular protein that can physically interact with the ECM, other matricellular proteins, and cell receptors.^25^ Latent TGFβ is one of the growth factors reposited in the ECM.^35^ THBS-1 activates TGFβ by direct interactions with latent TGFβ in ECM.^7, 36^ Since TGFβ is a key driver of myofibroblast differentiation and ECM production, this THBS-1-mediated activation promotes fibrotic responses. Hsieh et al. demonstrated that THBS-1 is elevated in EoE fibroblasts by proteomic analysis and that it alters the function of normal fibroblasts.^37^ In this study, we have shown that *THBS-1* is decreased in remission fibroblasts compared with active EoE fibroblasts and that its genetic knock-down reduces ECM production. This result suggests that reducing THBS-1 may assist in myofibroblast dedifferentiation by limiting latent TGFβ activation. Of note, *THBS-1* knockdown did not alter αSMA expression, which may be due to the limitations of performing these experiments in vitro or may suggest the effect is related more to production of ECM. Nevertheless, since THBS-1 and TGFβ create a feed-forward loop in fibrostenosis, THBS-1 downregulation may be indirectly contributing to myofibroblast dedifferentiation by limiting the activation of latent TGFβ and breaking the cycle.

Transcriptome analyses on TGFβ stimulated fibroblasts and active EoE fibroblasts identified the Integrin 1 pathway as one of the most enriched pathways. Integrins are transmembrane receptors that mediate the adhesion of cells to the ECM or neighboring cells. Integrin-mediated signaling allows cells to regulate crucial biological processes, including survival, proliferation, and differentiation.^26, 33, 38^ Given that THBS-1 has an interaction for integrin receptors^39–41^, its reduction resulting from PGE₂ signaling could downregulate the integrin pathway and thereby suppress fibroblast proliferation and latent TGFβ activation.

Currently, integrin antagonists have indications for inflammatory conditions including inflammatory bowel disease and multiple sclerosis.^42^ Evaluating this pathway is warranted to examine how integrin-directed therapy may provide anti-inflammatory and anti-fibrotic effects for conditions such as EoE.

One weakness of this study is that we limited out focus to fibroblasts. How PGE₂ may affect the esophageal epithelium or its inflammatory infiltrate in EoE is unknown.

Another limiting factor is our use of tissue-culture plastic, an extremely common culture substrate despite it being a potent fibroblast activator.^5^ Despite the culture conditions, we found PGE₂ induced myofibroblast dedifferentiation. The results observed in the EoE mouse model also support the effectiveness of PGE₂ in suppressing fibrostenosis.

In conclusion, we demonstrated that PGE₂ dedifferentiates esophageal myofibroblasts via the cAMP/YAP/THBS-1 pathway. Targeting PGE₂, its receptor, or downstream targets may be a new therapeutic strategy for severe EoE with esophageal stenosis. Furthermore, elucidating these mechanisms may have implications for fibrostenosis beyond EoE, including gastroesophageal reflux disease and anastomotic stricture after esophageal resection or esophageal atresia repair.

## Acknowledgments

CHOP Gastrointestinal Epithelium Modeling Program and Core (RRID: SCR_026402). The Center for Molecular Studies in Digestive and Liver Diseases (RRID: SCR_022420).

## Abbreviations

AC: (Adenylyl Cyclase)
ANOVA: (Analysis of Variance)
COL1A1: (Collagen Type I Alpha 1 Chain)
DAPI: (4′,6-Diamidino-2-Phenylindole)
DEGs: (Differentially Expressed Genes)
EoE: (Eosinophilic Esophagitis)
ECM: (Extracellular Matrix)
EPR: (E-prostanoid G protein-coupled receptors)
eos: (Eosinophils)
FDR: (False Discovery Rate)
FEF3: (fetal esophageal fibroblast)
FN1: (Fibronectin 1)
GAPDH: (glyceraldehyde-3-phosphate dehydrogenase)
GSEA: (Gene Set Enrichment Analysis)
hpf: (High Power Field)
IP: (Immunoprecipitation)
IPF: (Idiopathic Pulmonary Fibrosis)
NES: (Normalized Enrichment Score)
PGE₂: (Prostaglandin E₂)
PEF: (Patient-derived esophageal fibroblast)
PKA: (Protein Kinase A)
SD: (Standard Deviation)
αSMA: (Alpha–Smooth Muscle Actin)
TGFβ: (Transforming Growth Factor Beta)
THBS-1: (Thrombospondin-1)
YAP: (Yes-Associated Protein)

---

**Supplementary Figure 1.**
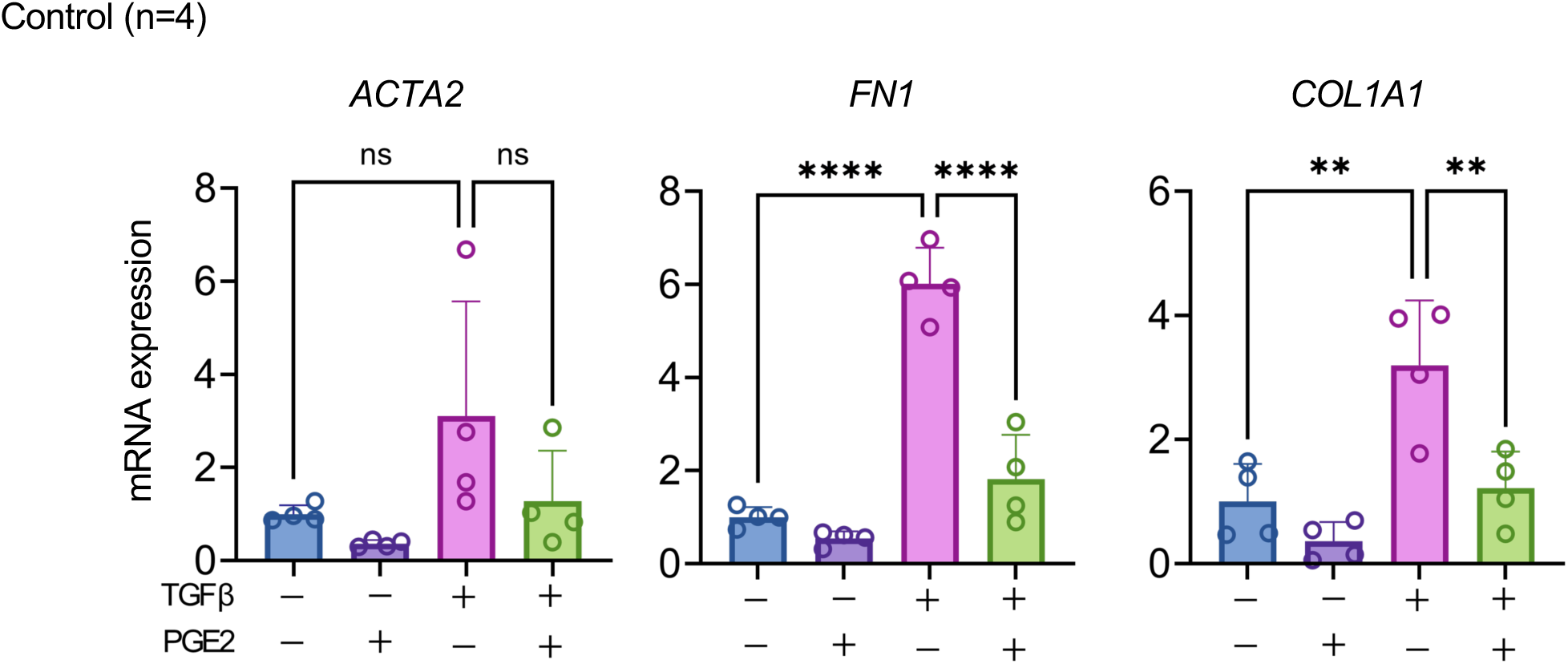
Relative *ACTA2*, *FN1*, and *COL1A1* expression by qPCR in control PEF stimulated with TGFβ with or without PGE2 (1 μM) for 24 hours (n = 4). 1-way analysis of variance was performed for statistical analyses. **p < 0.01, ****p < 0.0001, ns not significant

**Supplementary Figure 2.**
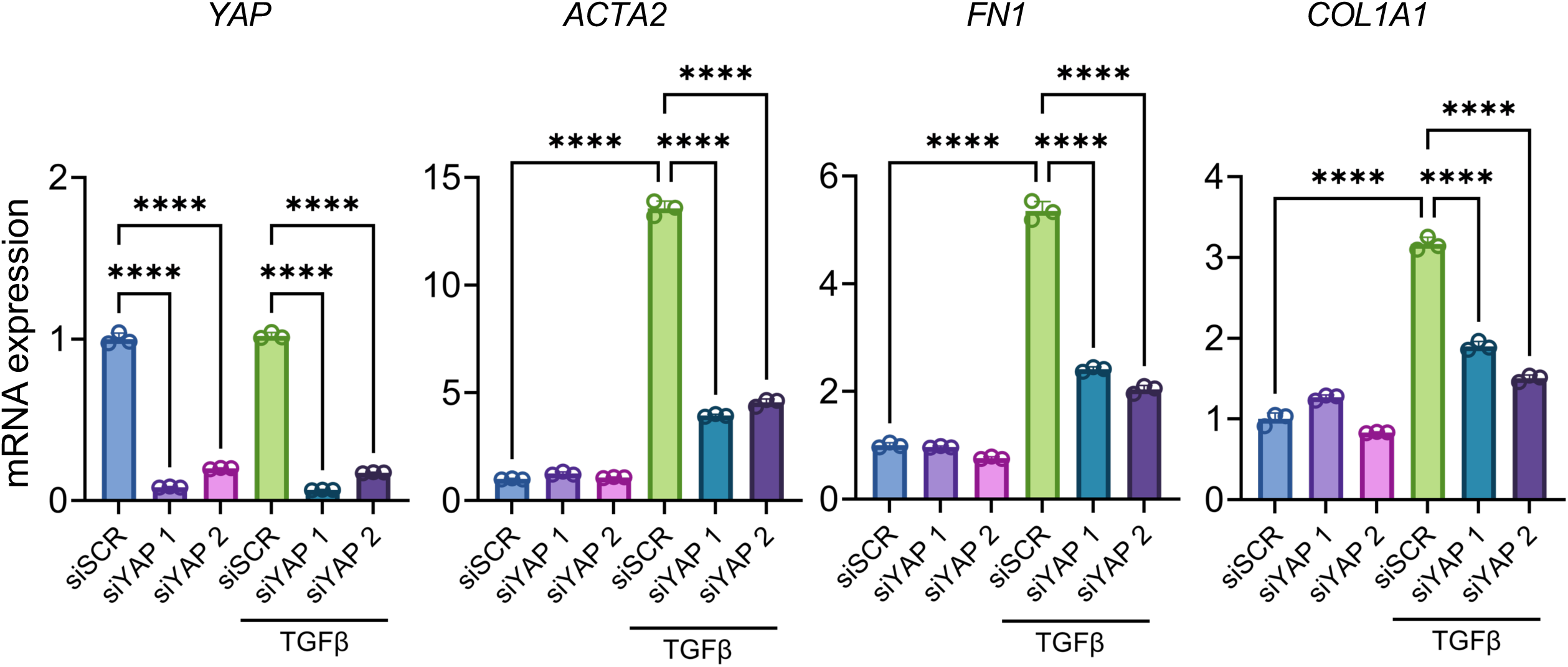
Transfection with siRNA against YAP in FEF3 cells. Scramble siRNA-transfected FEF3 cells were used as a control. Twenty-four hours after transfection, FEF3 was stimulated with TGFβ. Relative *YAP*, *αSMA*, *FN1*, and *COL1A1* gene expression by qPCR.

**Supplementary Figure 3.**
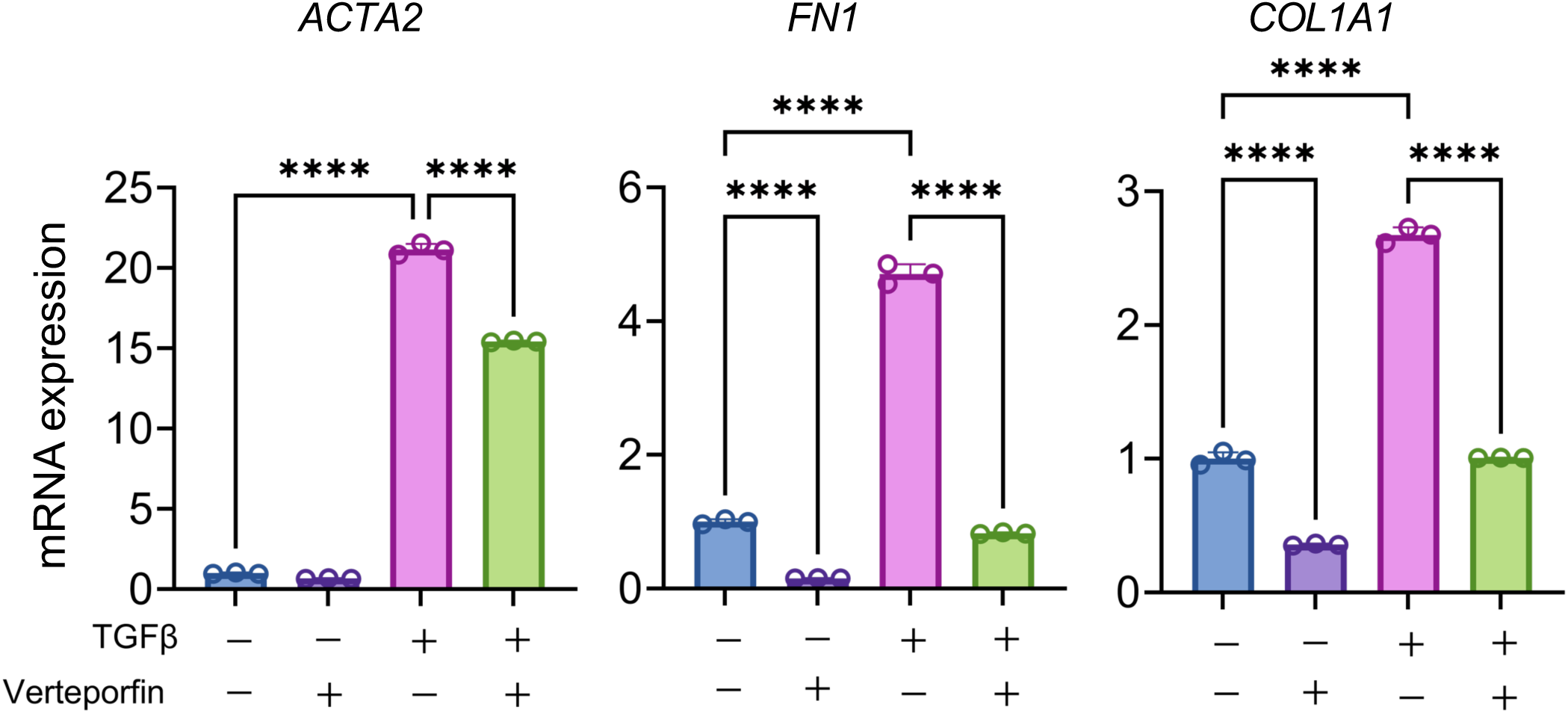
Relative *ACTA2*, *FN1*, and *COL1A1* expression in FEF3 stimulated with TGFβ with or without verteporfin (1 μM) for 24 hours (n = 3).

**Supplementary Figure 4.**
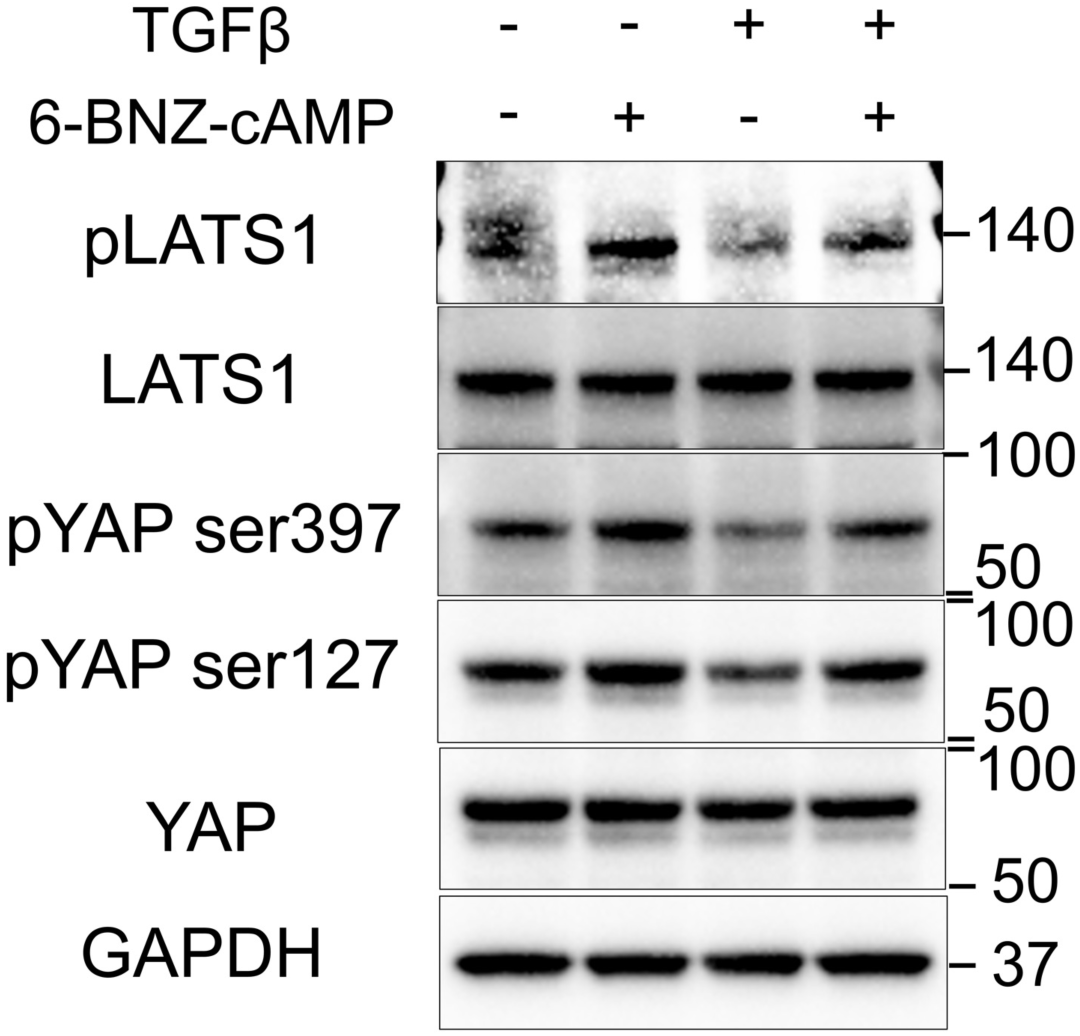
Representative Western blots showing phosphorylation of LATS and YAP were determined 30 minutes after treatment with TGFβ with or without 6BNZ-cAMP (1 mM), respectively.

**Supplementary Figure 5.**
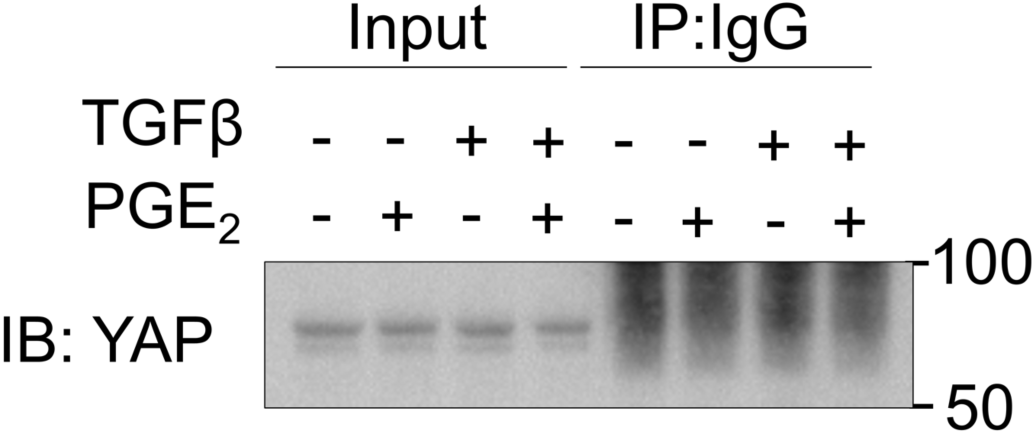
Negative control for immunoprecipitation. Cell lysates were immunoprecipitated with control IgG, and the precipitates were analyzed by immunoblotting with YAP.

**Supplementary Figure 6.**
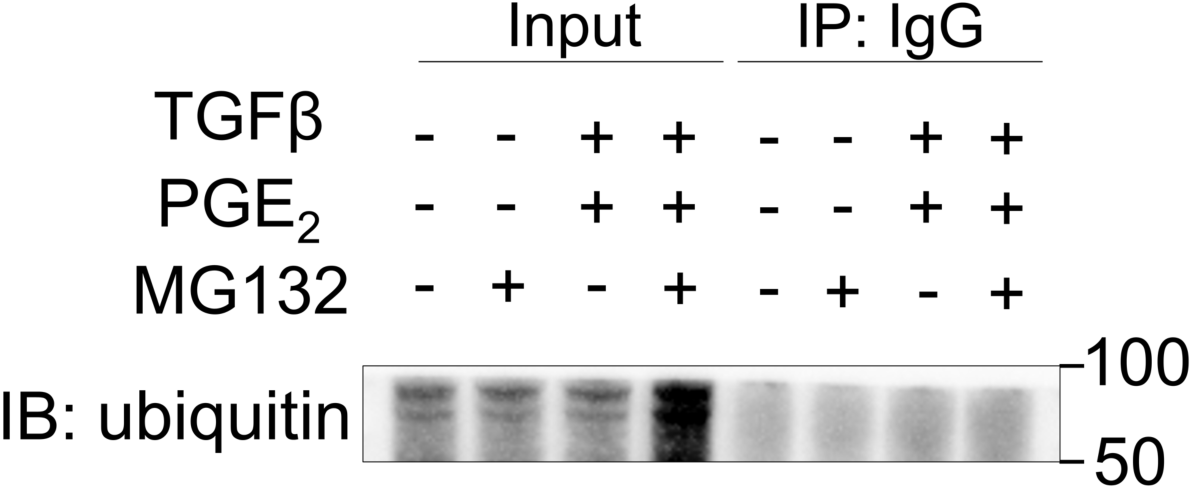
Negative control for immunoprecipitation. Cell lysates were immunoprecipitated with control IgG, and the precipitates were analyzed by immunoblotting with ubiquitin.

**Supplementary Figure 7.**
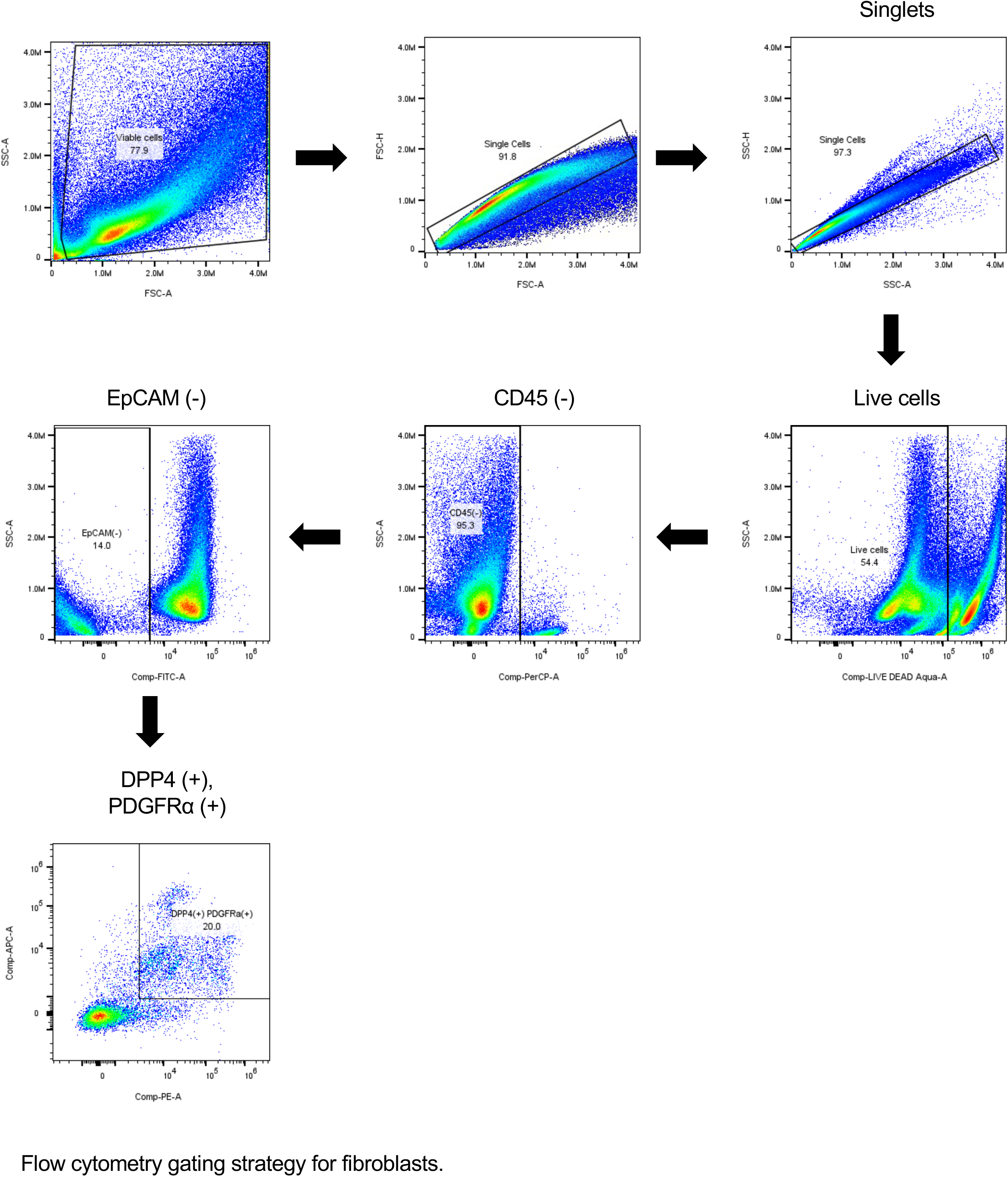
Flow cytometry gating strategy for fibroblasts.

**Supplemental Table S1.** Patient demographics in this study.

| **ID** | **Application** | **Age (years)** | **Gender** | **Diagnosis** | **Max Eos/hpf** | **History of impaction** | **History of stricture** | **Symptom of dysphagia** | **Symptom of regurgitation** |
| --- | --- | --- | --- | --- | --- | --- | --- | --- | --- |
| 1 | Fig 1E PCR | 43.0 | Male | Active-EoE | 50 | No | No | Yes | Yes |
| 2 | Fig 1E PCR | 46.0 | Male | Active-EoE | 30 | Yes | Yes | Yes | Yes |
| 3 | Fig 1E PCR | 21.0 | Female | Active-EoE | 17 | No | No | Yes | No |
| 4 | Fig 1E PCR | 43.0 | Female | Active-EoE | 100 | No | No | No | No |
| 5 | Fig 1E PCR | 43.0 | Female | Active-EoE | 35 | No | No | No | No |
| 6 | Fig 1E PCR | 17.3 | Male | Active-EoE | 40 | No | No | No | No |
| 7 | Fig 1E PCR | 10.7 | Male | Active-EoE | 80 | No | No | No | No |
| 8 | Fig 1E PCR | 10.5 | Male | Active-EoE | 50 | No | No | No | No |
| 9 | Fig 1E PCR | 11.0 | Male | Active-EoE | 70 | No | No | No | No |
| 10 | Fig 1E PCR | 13.6 | Male | Active-EoE | 50 | No | No | Yes | Yes |
| 11 | Fig 1E PCR | 13.1 | Male | Active-EoE | >50 | No | No | No | No |
| 12 | Fig 1E PCR | 6.9 | Female | Active-EoE | 25 | No | No | Yes | Yes |
| 13 | Fig 1E PCR | 14.1 | Male | Active-EoE | >50 | No | Yes | No | No |
| 14 | Fig 3A Fig5D IHC | 6.6 | Male | Active-EoE | 65 | No | No | Yes | No |
| 15 | Fig 3A Fig5D IHC | 11.4 | Male | Active-EoE | >50 | Yes | No | Yes | No |
| 16 | Fig 3A Fig5D IHC | 6.3 | Male | Active-EoE | 30-40 | Yes | No | No | No |
| 17 | Fig 3A Fig5D IHC | 11.1 | Male | Active-EoE | 50 | Yes | No | Yes | No |
| 18 | Fig 3A Fig5D IHC | 6.7 | Male | Active-EoE | >100 | Yes | No | Yes | Yes |
| 19 | Fig 3A Fig5D IHC | 8.7 | Female | Control | 0 | No | No | No | No |
| 20 | Fig 3A Fig5D IHC | 9.5 | Male | Control | 0 | No | No | No | Yes |
| 21 | Fig 3A Fig5D IHC | 8.2 | Female | Control | 0 | No | No | No | No |
| 22 | Fig 3A Fig5D IHC | 8.5 | Male | Control | 0 | No | No | No | No |
| 23 | Fig 3A Fig5D IHC | 9.1 | Male | Control | 1 | No | No | No | No |
| 24 | Fig 3A Fig5D IHC | 12.7 | Male | Remission | 1 | mild symptoms of food impaction | No | Yes | No |
| 25 | Fig 3A Fig5D IHC | 15.9 | Male | Remission | 1 | No | No | No | No |
| 26 | Fig 3A Fig5D IHC | 15.3 | Male | Remission | 10 | No | No | No | No |
| 27 | Fig5B PCR | 10.5 | Male | Active-EoE | 50 | No | No | No | No |
| 28 | Fig5B PCR | 11.0 | Male | Active-EoE | 70 | No | No | No | No |
| 29 | Fig5B PCR | 13.6 | Male | Active-EoE | 50 | No | No | Yes | Yes? |
| 30 | Fig5B PCR | 6.1 | Male | Active-EoE | 80 | No | No | No | No |
| 31 | Fig5B PCR | 17.7 | Male | Active-EoE | 62 | Yes | Yes | Yes | No |
| 32 | Fig5B PCR | 12.3 | Male | Active-EoE | 20 | No | No | No | No |
| 33 | Fig5B PCR | 10.8 | Male | Active-EoE | 18 | No | No | No | Yes |
| 34 | Fig5B PCR | 11.7 | Male | Remission | 2 | Yes | No | Yes | No |
| 35 | Fig5B PCR | 10.7 | Male | Remission | 0 | no official dx but has complained of food impaction in the past | No | Yes | Yes |
| 36 | Fig5B PCR | 6.1 | Male | Remission | unremarkable<br>squamous<br>mucosa | No | No | Yes | No |
| 37 | Fig5B PCR | 8.1 | Male | Remission | 8 | No | No | Yes | No |
| 38 | Fig5B PCR | 4.0 | Male | Remission | 1 | No | No | No | Yes |
| 39 | Fig5B PCR | 14.3 | Male | Remission | 5 | No | No | No | Yes |
| 40 | Fig5B PCR | 16.2 | Male | Remission | 0 | No | No | No | No |
| 41 | Supplemental<br>Figure 1 PCR | 6.5 | Female | Control | 0 | No | No | Yes | No |
| 42 | Supplemental<br>Figure 1 PCR | 15.0 | Male | Control | 10 | No | No | No | No |
| 43 | Supplemental<br>Figure 1 PCR | 8.7 | Female | Control | 0 | No | No | No | No |
| 44 | Supplemental<br>Figure 1 PCR | 14.1 | Male | Control | 0 | No | No | No | No |

Eos, eosinophil; HPF, high-power field; PCR, polymerase chain reaction; IHC, immunohistochemistry

**Supplemental Table S2.**
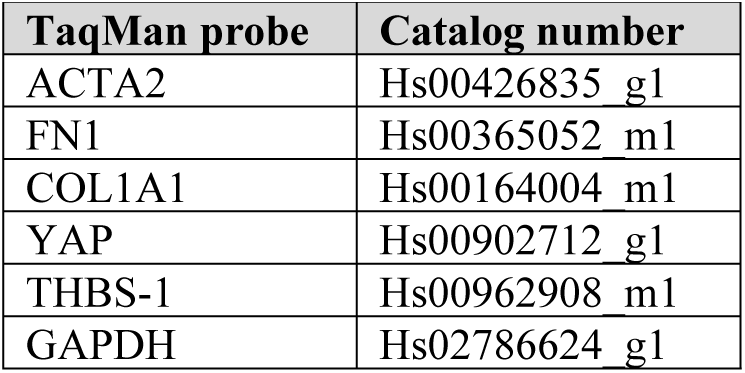
List of TaqMan probes used for qRT-PCR.

**Supplemental Table 3.** List of antibodies used.

| Antibody information |  |  |  | Dilution / Application |  |  |  |
| --- | --- | --- | --- | --- | --- | --- | --- |
| Antibody | Company | Catalog number | Description | WB | IP | ICC | IHC |
| Anti-αSMA | Cell Signaling Technology | 19245S | Rabbit monoclonal | 1:1000 | | | |
| Anti-α Smooth Muscle Actin | Sigma-Aldrich | A5228 | Mouse monoclonal | | | 1:500 | |
| Anti-FN1 | Cell Signaling Technology | 26836S | Rabbit monoclonal | 1:1000 |  | 1:1000 |  |
| Anti-COL1A1 | Cell Signaling Technology | 72026S | Rabbit monoclonal | 1:1000 |  | 1:1000 |  |
| Anti-phospho-LATS1 | Cell Signaling Technology | 8654S | Rabbit monoclonal | 1:500 |  |  |  |
| LATS1 | Cell Signaling Technology | 3477S | Rabbit monoclonal | 1:1000 |  |  |  |
| Anti-phospho YAP(Ser127) | Cell Signaling Technology | 13008S | Rabbit monoclonal | 1:1000 |  |  |  |
| Anti-phospho YAP(Ser397) | Cell Signaling Technology | 13619S | Rabbit monoclonal | 1:1000 | 1:100 |  |  |
| Anti-YAP | Cell Signaling Technology | 14074S | Rabbit polyclonal | 1:1000 |  | 1:1000 | Human 1:100, Mouse 1:50 |
| Anti-phospho-SMAD2 (Ser465/Ser467) | Cell Signaling Technology | 18338S | Rabbit monoclonal | 1:1000 |  |  |  |
| Anti-phospho SMAD3 (Ser423/425) | Cell Signaling Technology | 9520S | Rabbit monoclonal | 1:500 | 1:50 |  |  |
| Anti-Smad2/3 | Cell Signaling Technology | 8685S | Rabbit monoclonal | 1:1000 |  |  |  |
| Anti-Thrombospondin-1 | Cell Signaling Technology | 37879S | Rabbit monoclonal | 1:1000 |  |  |  |
| Anti-Thrombospondin-1 | Abcam | ab1823 | Mouse monoclonal |  |  |  | Human 1:20 |
| Anti-Thrombospondin-1 | Thermo Scientific | MA5-13398 | Mouse monoclonal |  |  |  | Mouse 1:25 |
| Anti-GAPDH | Thermo Fisher Scientific | MA5-15738 | Mouse monoclonal | 1:1000 |  |  |  |

WB, Western blot; IP, Immunoprecipitation; ICC, Immunocytochemistry; IHC, Immunohistochemistry

**Supplemental Table S4.** List of antibodies used.

| Antibody information |  |  |  | Dilution / Application |
| --- | --- | --- | --- | --- |
| Antibody | Company | Catalog number | Description | FC |
| Anti-CD45 | BioLegend | 103129 | Rat monoclonal | 1:100 |
| Anti-CD326 | BioLegend | 118208 | Rat monoclonal | 1:100 |
| Anti-CD26 | BioLegend | 137804 | Rat monoclonal | 1:20 |
| Anti-CD140a | BD Biosciences | 135908 | Rat monoclonal | 1:20 |
| Anti-Thrombospondin-1 | Thermo Scientific | MA5-13398 | Mouse monoclonal | 1:20 |
| Anti-α-smooth muscle actin antibody | Novus Biologicals | NBP2-34522AF405 | Mouse monoclonal | 1:100 |
| And anti-FN1 antibody | Novus Biologicals | NBP2-34504PECY7 | Mouse monoclonal | 1:100 |

FC, Flow cytometry

